# A multiomics comparison of weight loss interventions shows the distinct effects of bariatric surgery and intermittent fasting on hepatic metabolism

**DOI:** 10.1101/2022.06.13.495926

**Authors:** Arnon Haran, Michael Bergel, Doron Kleiman, Liron Hefetz, Hadar Israeli, Sarah Weksler-Zangen, Bella Agranovich, Ifat Abramovich, Rachel Ben-Haroush Schyr, Eyal Gottlieb, Danny Ben-Zvi

## Abstract

The prevalence of obesity and non-alcoholic fatty liver (NAFL) is steadily increasing. Weight loss can lead to improvement in NAFL in patients and in model organisms. Weight loss can be achieved through either dietary and lifestyle interventions, pharmacotherapy, and bariatric surgery. Some interventions, such as intermittent fasting, have been purported to have benefits beyond those conferred by weight loss alone.

In this study, rats were provided a high-fat, high-sucrose diet for 7 weeks and then were assigned to strict caloric restriction with intermittent fasting (IF-CR) or to sleeve gastrectomy (SG) bariatric surgery, achieving equivalent weight loss. The two interventions were compared to ad-libitum fed controls. The metabolome of the blood entering the liver was analyzed in the three experimental groups, as well as the metabolome and transcriptome of the liver under fasting conditions, to test how different means of weight loss affect the input of the liver and hepatic metabolism.

Surprisingly, the two interventions had different and often opposite effects on the composition of portal blood and liver metabolites. IF-CR resulted in sweeping changes across multiple central metabolic pathways, including an increase in de novo lipogenesis and triglyceride export, and upregulation of glycolysis, citric acid cycle, and the pentose phosphate pathway, as well as an increase in glycogen storage. Meanwhile, the effects of SG concentrated on one-carbon metabolic pathways, with changes in metabolite and transcript levels pointing to decreased transmethylation and methionine cycle activity, with downstream effects on levels of glutathione and the hepatic oxidative state.

In conclusion, our study highlights how different means of weight loss affect distinct metabolic pathways and demonstrates a unique effect of bariatric surgery on hepatic one-carbon and redox pathways.

## Introduction

The worldwide prevalence of obesity is steadily increasing. In 2015, an estimated 12.0% of adults and 5.0% of children worldwide had obesity. Excess body weight accounted for about 4 million deaths and 120 million disability-adjusted life-years, most of these due to cardiovascular disease and diabetes^1^. Weight loss can prevent many of the adverse consequences of obesity, and can be achieved through either dietary and lifestyle interventions, pharmacotherapy, or bariatric surgery^2^. Some of these interventions have been purported to have benefits beyond those conferred by weight loss alone. For example, with regards to mortality and cardiovascular adverse events, the benefits of bariatric surgery were evident even with minimal weight loss, and greater degrees of weight loss were required to achieve the same effects through dietary intervention alone^3,4^. Likewise, an intermittent fasting (IF) regimen resulted in improved insulin sensitivity when compared to a similar degree of weight loss achieved through continuous energy restriction diets, both in humans and in animal models^5–7^, in addition to reported benefits for unrelated morbidities such as multiple sclerosis or cancer^8^. The mechanisms underlying these beneficial effects remain largely unknown. For both bariatric surgery and IF, numerous metabolic alterations are observed following the intervention, which are not seen in weight-matched controls. These include changes in serum amino acid and bile acid concentrations^9,10^ and in the composition of the gut microbiome^11,12^. However, the clinical significance of these alterations has been questioned^13,14^. We have previously shown a specific and weight loss-independent effect of bariatric surgery on hepatic insulin sensitivity^15,16^, suggesting that changes in liver metabolism may explain some of the positive effects of surgery. The liver is a key organ in the fasting response, and mediates many of the beneficial effects of IF as well^7^.

Obesity is associated with an alteration of hepatic fatty acid (FA) metabolism leading to an excessive accumulation of lipids within hepatocytes, a condition known as non-alcoholic fatty liver (NAFL) or hepatic steatosis^17,18^. The close link between obesity and hepatic steatosis is demonstrated by an increase in NAFL prevalence with increasing body mass index^19^. From a metabolic standpoint, NAFL may be understood as a condition in which the liver’s capacity to handle FAs and other metabolic energy substrates is overwhelmed^20^.

Free FAs in the liver are derived either from dietary lipids, from adipose tissue through mobilization of triglyceride (TG) stores, or from hepatic de novo lipogenesis (DNL). Within hepatocytes, FAs are ultimately either oxidized or converted to TGs, which are then exported as very-low-density lipoprotein (VLDL) particles or stored intrahepatically. Thus, lipid accumulation in hepatocytes may be due to either excess FA influx or an impaired ability to dispose of FAs through ß-oxidation or TG export^17^.

FA metabolism requires the contribution of several other central metabolic pathways in the liver. DNL hinges on a supply of acetyl-CoA and reducing power in the form of NADPH, derived from the pentose phosphate pathway, the citric acid cycle, and serine metabolism^21^. Likewise, VLDL assembly and TG export rely on the liver’s ability to synthesize sufficient phosphatidylcholine (PC), either directly from dietary choline, or via the methylation of phosphatidylethanolamine (PE) in a reaction catalyzed by phosphatidylethanolamine N-methyltransferase (PEMT). Fat export is therefore dependent on the availability of methyl group donors.

The conversion of PE to PC is one of the two major methyl-consuming reactions in the liver and the body as a whole^22,23^, along with the methylation of guanidinoacetic acid (GAA) to produce creatine. Methyl groups are also required for nucleotide synthesis, maintenance of DNA methylation, protein and RNA methylation, and numerous other reactions. Methyl transfer reactions utilize S-adenosylmethionine (SAM) as the universal methyl donor, which is converted to S-adenosylhomocysteine (SAH) in the process. Methionine is converted to SAM and thereby enters the methionine cycle in a reaction catalyzed by methionine adenosyltransferases (MATs). Three MAT isoenzymes exist, of which MAT I and MAT III, both encoded by the *MAT1A* gene, predominate in the liver^24^. Regeneration of SAM involves the transfer of methyl groups to homocysteine via either choline-derived methyl donors such as betaine, or via folic acid and vitamin B12-dependent pathways, utilizing methyl groups generated from the metabolism of, e.g., serine or glycine. Depletion of hepatic SAM, by way of a diet low in choline and methionine^25^ results in decreased PC and fat accumulation within the liver due to impaired VLDL export capacity^26^. Homocysteine generated via the methionine cycle is also a precursor for synthesis of the antioxidant glutathione. Therefore, decreased entry of methionine into the methionine cycle is expected to result not only in depletion of SAM but also decreased production of glutathione, as indeed seen in some cases with methionine restricted diets^27,28^ and, more directly, with knockout or silencing of *MAT1A*^29^. An excess of SAM is as injurious as its depletion, and may lead to the development of hepatic steatosis^30,31^. Likewise, in different settings, knockout or silencing of the *MAT1A* gene has been seen to either result in increased susceptibility to hepatic steatosis^24^ or protect from diet-induced NAFL^29^.

In the present work, we seek to gain insight into which of the above-mentioned metabolic pathways are affected by weight loss interventions, and whether different interventions differ in this regard. We analyzed the metabolome of the hepatic blood supply and the metabolome and transcriptome of the liver itself following sleeve gastrectomy (SG), a common bariatric surgery, and IF with caloric restriction (IF-CR), in rats fed a high-fat, high-sucrose diet. Both interventions led to a similar loss of weight but resulted in distinct and in some cases contradictory effects on several metabolic pathways. The effect of bariatric surgery is apparent chiefly within pathways related to methionine and methyl group metabolism. In contrast, IF-CR affects primarily fatty acid metabolism and synthesis, which remain relatively unaffected by SG.

## Methods

### Experimental model: animals, diet, and surgical procedures

All experiments were approved by the institutional animal care and use committee. Male Sprague-Dawley rats were used for the studies. Animals were housed in pairs and were maintained on a 12-hour light, 12-hour dark cycle.

The animals were fed a high-fat, high-sucrose diet (Envigo TD.08811; HFHSD) for a total of 12 weeks, during which they were divided into three experimental groups (figure 1A). One group (8 rats) underwent sleeve gastrectomy (SG) following seven weeks of high-fat, high-sucrose feeding, a second group (6 rats) underwent sham operation at the same time and then underwent an IF-CR diet to match the weight of the SG-treated rats, and a third group (6 rats) was allowed to consume food freely and did not undergo any intervention (*ad libitum* [AL]). During caloric restriction, food was provided once daily in the evening. The amount of food was initially restricted to 10 grams/animal/day to induce weight loss similar to that of the SG group and was then gradually increased up to 20 grams/animal/day as both groups gained weight in parallel.

**Figure 1:**
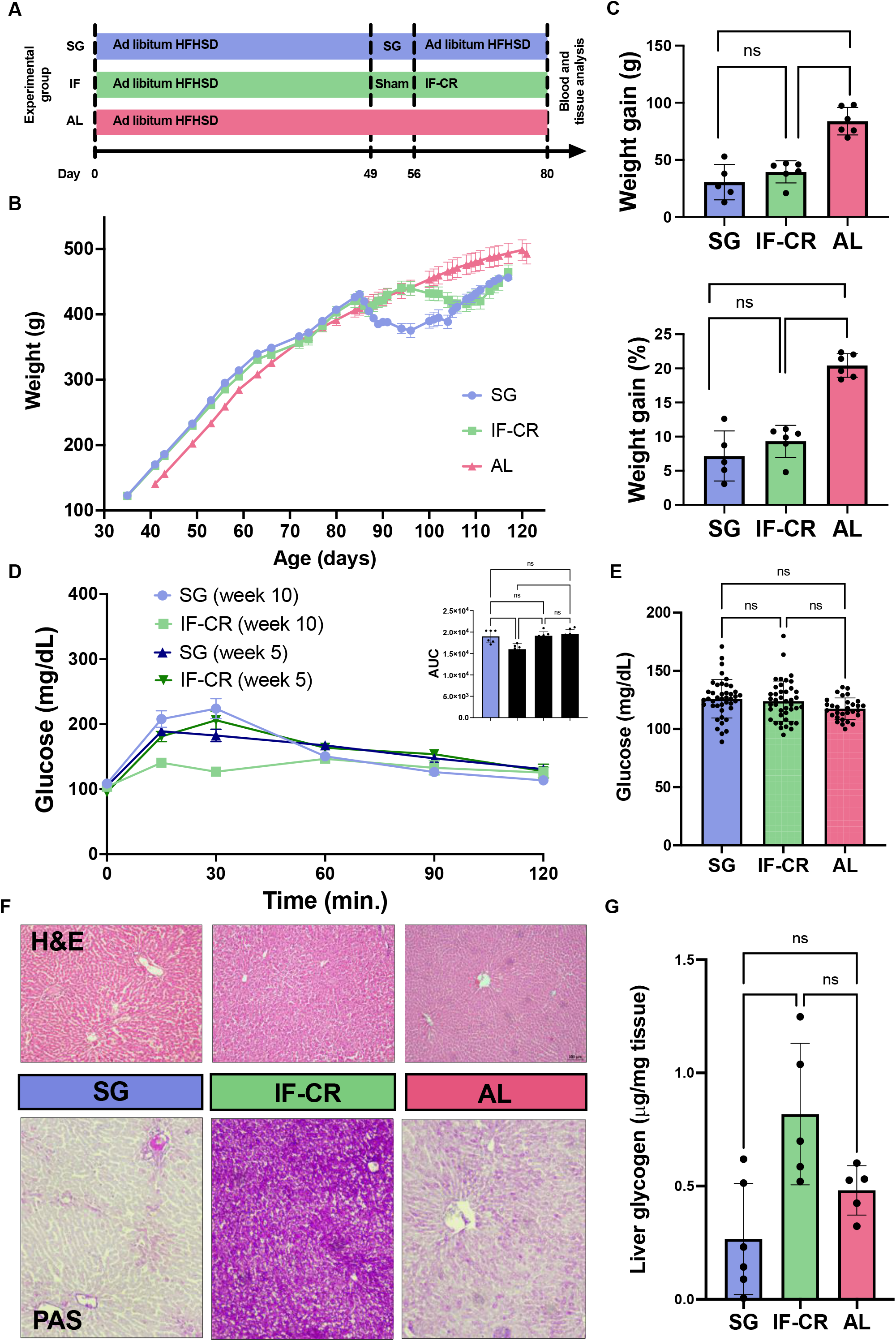
Overview and liver histology. (A) Experimental design. Animals were fed a high-fat, high-sucrose diet (HFHSD) for 7 weeks, and then underwent either sleeve gastrectomy (SG), sham surgery followed by intermittent fasting and caloric restriction (IF-Cr), or no intervention (AL). At day 80, animals were sacrificed and blood and tissue collected for analysis. (B) Body weight of animals in the 3 experimental groups (mean +/-SEM). (C) Weight gain from day 49 (pre-intervention) to day 80 in the 3 experimental groups, in grams (top) and as a % of day 49 weight (bottom). (D) Oral glucose tolerance tests carried out at week 5 (pre intervention) and week 10 (post intervention) in the SG and IF-CR groups. Inset: area under the curve (AUC). (E) Morning non-fasting serum glucose measurements carried out in the three experimental groups. (F) Top: hematoxylin and eosin-stained liver sections, 10X magnification. Bottom: periodic acid-Schiff-stained liver sections, 10X magnification. (G) Glycogen content in liver homogenates. ** p<0.01 ANOVA with Tukey post-hoc test.

Rats were fasted for the 12 hours before and for the first 24 hours after surgery and were then maintained on a liquid diet for a further 72 hours, following which the preoperative diet was returned. Surgery was performed under inhalational anesthesia (isoflurane) and post-operative analgesia (meloxicam) was administered daily for the 5 days following surgery.

SG was performed with the use of a Covidien Endo GIA surgical stapler. A midline incision was made, and the stapler was positioned to achieve a removal of 70-80% of the stomach, following previously described protocols^32^. For sham operations, a midline incision was followed by manual manipulation of the stomach only.

Oral glucose tolerance tests (OGTT) were carried out in the SG and IF-CR groups at week 5 (pre intervention) and week 10 (post intervention). Oral gavage of 2 g/kg glucose was administered following an overnight fast, and serum glucose was determined using an Accu-Chek Performa glucometer (Roche Diagnostics Ltd.) at intervals of 15-30 minutes up to 120 minutes post-gavage. Random, non-fasting measurements of serum glucose were also carried out in all experimental groups once to twice weekly.

At week 12, five weeks after surgery in the interventional groups, and following a 10-14-hour overnight fast, blood samples from the portal vein and the aorta were obtained, as well as tissue samples from the liver. Anesthesia was achieved with intraperitoneal phenobarbital. All samples were immediately frozen in liquid nitrogen and then stored at -80°C until their analysis.

All animals in the IF-CR and AL groups were included in the final analysis. In the SG groups, two animals died due to surgical complications and one animal died prior to obtaining blood and tissue samples due to anesthetic complications. Therefore, data from only five animals in this group were included in the final analysis.

### Analysis

Metabolomic analysis of the hepatic input and metabolomic and transcriptomic analysis of the liver itself were performed in the three experimental groups. Portal and aortic blood samples were analyzed separately, and the hepatic input was then defined as their combination in a 4:1 ratio^33^.

Metabolite extraction was performed with a methanol/acetonitrile (3:1) mixture. Quality control standards and a quality control mix of all samples were used to assess for technical variability. Samples were then analyzed using HPLC/MS and raw data were extracted and matched to a reference library using the software TraceFinder (Thermo Scientific) and based on expected retention time and mass. Automatic identification was followed by manual curation to ensure its accuracy. Peaks were quantified using the area under the curve and normalized to the total ion count measured for each sample. All metabolomic statistical analyses were carried out using MetaboAnalyst^34^.

RNA was isolated from liver tissue using a TRIzol reagent-based protocol. Quantity and quality of the extracted RNA were measured using the Agilent TapeStation system. RNA integrity numbers were >8.2 for all samples. cDNA libraries were prepared using approximately 1000 ng of RNA per sample and following poly(A) enrichment and were then sequenced on the Illumina NextSeq 500 system. After alignment to the reference genome, differential gene expression analysis was carried out using DESeq2 and Matlab. Low expression genes were filtered out prior to analysis based on a threshold of average expression of less than 1 copy per million across samples.

Phosphatidylcholine, phosphatidylethanolamine, and glycogen content in liver tissue were measured using Abcam colorimetric/fluorimetric assays (ab83377, ab241005, and ab65620, respectively), following the manufacturer’s instructions. Serum triglycerides, low-density lipoprotein (LDL) cholesterol, and high-density lipoprotein (HDL) cholesterol, were determined using a Cobas c111 analyzer (Roche Diagnostics Ltd.).

One-way analysis of variance (ANOVA) was used to recognize individual metabolites or genes whose concentrations or expression, respectively, were significantly altered between the experimental groups. Correction for multiple comparisons was carried out using the Benjamini-Hochberg procedure^35^ with a false discovery rate (FDR) or adjusted p-value cutoff of 0.05. ANOVA post-hoc analyses were performed using Fisher’s least significant differences (LSD) procedure.

Multivariate dimensionality reduction using partial least squares-discriminant analysis (PLS-DA) was used to identify groups of metabolites discriminating between the experimental groups. Metabolites were defined as belonging to components one, two, or neither, based on their variable importance in projection (VIP) score for that component being above 1.0 (an accepted threshold^36^) and greater than the VIP score for the other component. Pathway enrichment of each component was then carried out using the small molecule pathway database (SBPDB).

For differentially expressed genes, enrichment analysis by KEGG pathway was performed separately for upregulated and downregulated genes in each of the intervention groups (i.e., SG and IF-CR); these were defined as genes identified by ANOVA and post-hoc analysis as significantly altered against at least one of the other experimental groups, and with either a negative or positive fold-change compared to both other groups.

## Results

At the conclusion of 12 weeks of HFHSD, animals attained an average weight of 456.3 g (+/-14.4) in the SG group, 464.7 (+/-26.6) in the IF-CR group, and 492.8 (+/-39.2) in the AL group (figure 1B). Average weight gain after the intervention was 30.6 g (7.2%), 39.5 g (9.3%), and 83.9 g (20.4%) in the SG, IF-CR, and AL groups, respectively (figure 1C). OGTT results were not significantly affected by SG (figure 1D), whereas the IF-CR group showed improved glucose tolerance following the intervention, as manifested in a decreased glucose area under the curve. Morning non-fasting serum glucose levels were unaltered between the experimental groups (figure 1E).

Livers were examined histologically, which revealed a relative abundance of glycogen in the IF-CR group (figure 1F). Other than this finding, which was corroborated by a biochemical glycogen assay (figure 1G), there were no significant differences in liver histology, which appeared normal.

### Metabolomic analysis of serum entering the liver

In analysis of the metabolic composition of the hepatic input, 87 of 197 metabolites were identified as altered between the experimental groups (figure 2A-C). PLS-DA showed separation of the IF-CR group from the SG and AL groups along the first component, and of the SG group from the two other groups primarily along the second component (figure 2D). Pathway enrichment analysis showed component one to be enriched for fatty acid beta oxidation and biosynthesis, linolenic and linoleic acid metabolism, and ketone body metabolism (figure 2E). Component two was enriched for glycine and serine metabolism, glutathione metabolism, carnitine synthesis, and methionine metabolism (figure 2F).

**Figure 2:**
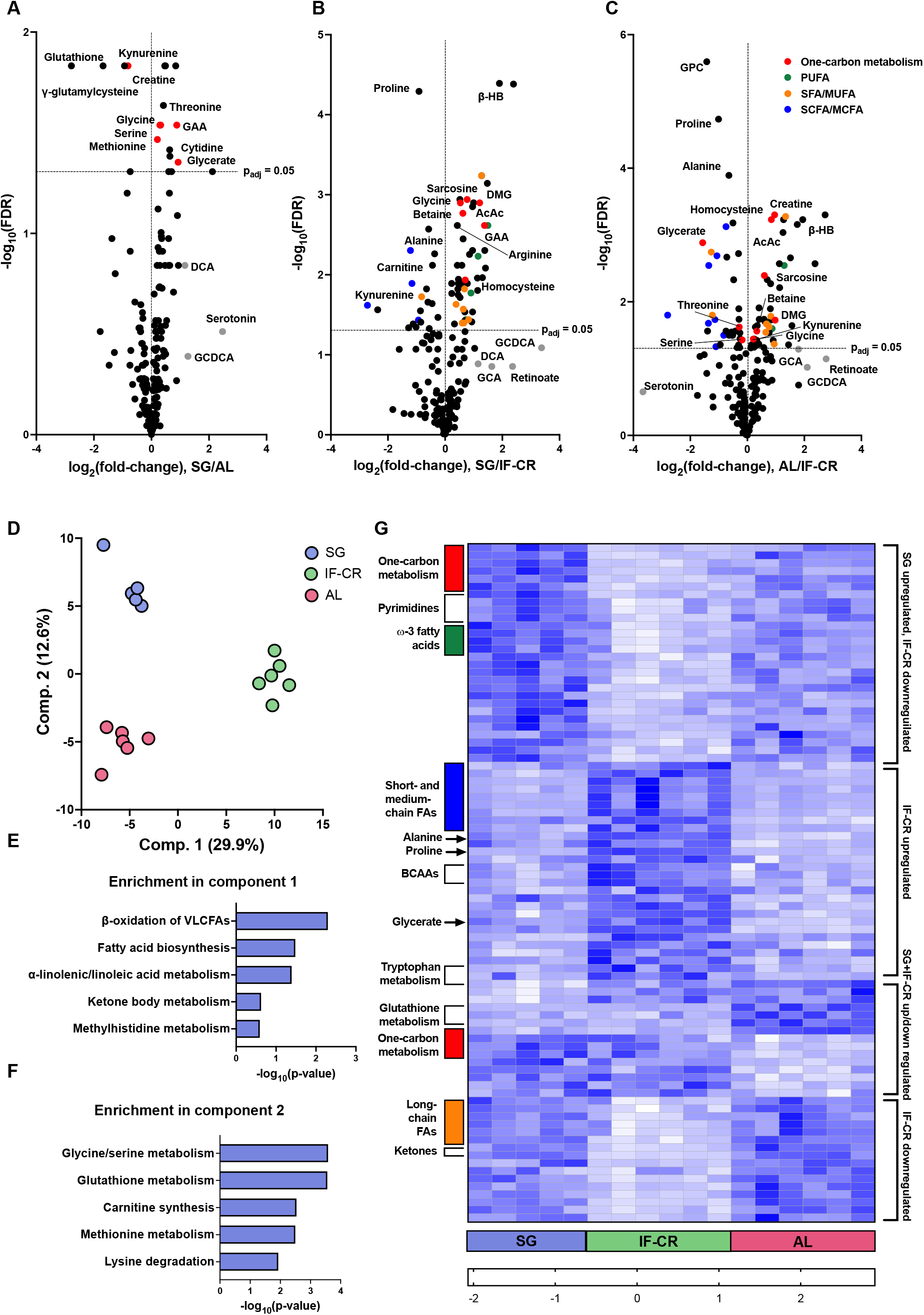
Metabolomic analysis of blood entering the liver. (A-C) Volcano plots showing the relative changes in metabolite concentrations in the hepatic input (80% portal blood, 20% aortic blood) between the experimental groups (SG: sleeve gastrectomy; IF-CR: intermittent fasting caloric restriction; AL: ad libitum; FDR = false discovery rate-adjusted p-value of between-group t-tests; PUFA: poly-unsaturated long-chain fatty acids; SFA/MUFA: saturated/mono-unsaturated long-chain fatty acids; SCFA/MCFA: short-/medium-chain fatty acids). (D) Partial least squares-discriminant analysis (PLS-DA) scores plot of metabolite abundance demonstrating separation of the IF-CR group from the AL group along the first component and of the SG group from the AL group along the second component. (E-F) Enrichment analysis by small molecule pathway database (SBPDB) pathways of metabolites in components 1 (E) and 2 (F) of PLS-DA. Metabolites were defined as belonging to component one, two, or neither, based on their VIP score for that component being above 1.0 and greater than the VIP score for the other component. (G) Heatmap of 87 metabolites identified as significantly altered between the experimental groups by ANOVA with FDR of <0.05. Color-code represents log_2_ fold-change between each sample and the mean level of the metabolite.

Within the differentially regulated metabolites identified by ANOVA, several patterns of between-group differences were seen (figure 2G). One group of compounds showed opposite trends in the IF-CR and SG group when compared to the AL group, being upregulated in the SG group, and downregulated in the IF-CR group. Among these were numerous metabolites related to one-carbon metabolism and methyl transfer reactions, as detailed below. Other metabolites showing a similar pattern were the pyrimidine nucleotides (cytidine and thymidine) and the polyunsaturated ω-3 long-chain fatty acids (LCFAs).

Some metabolites were downregulated or upregulated in the IF-CR group when compared to the AL group, but were relatively unaffected by SG. Metabolites downregulated in the IF-CR group include the ketone bodies acetoacetate and hydroxybutyrate and certain saturated and mono-unsaturated LCFAs. Notable compounds upregulated with IF-CR include the gluconeogenic precursors alanine and glycerate, proline, the branched-chain amino acids valine, leucine, and isoleucine, and gut microbiota-derived metabolites, including short- and medium-chain fatty acids (butyric, caproic, and caprylic acids), and the tryptophan breakdown product indole 3-acetate (IAA), which functions as a central mediator in gut microbiota-host interactions^37,38^. The concentrations of kynurenine, another tryptophan breakdown product, were high in the AL group, intermediate in the IF-CR group, and markedly decreased in the SG group.

### Metabolomic analysis of liver

In the liver itself, significant changes were seen in the concentrations of 51 of 241 metabolites across the experimental groups (figure 3A-C). PLS-DA showed separation of the IF-CR group from the SG and AL groups along the first component, and separation of the SG group from the two other groups along the second component (figure 3D). Pathway enrichment showed component one to be enriched for glutamate metabolism, fatty acid and plasmalogen biosynthesis, purine metabolism, and carnitine synthesis. Component two was enriched for glycine and serine metabolism as well as glycolysis and pentose phosphate pathway-related metabolites and ammonia recycling (figure 3E).

**Figure 3:**
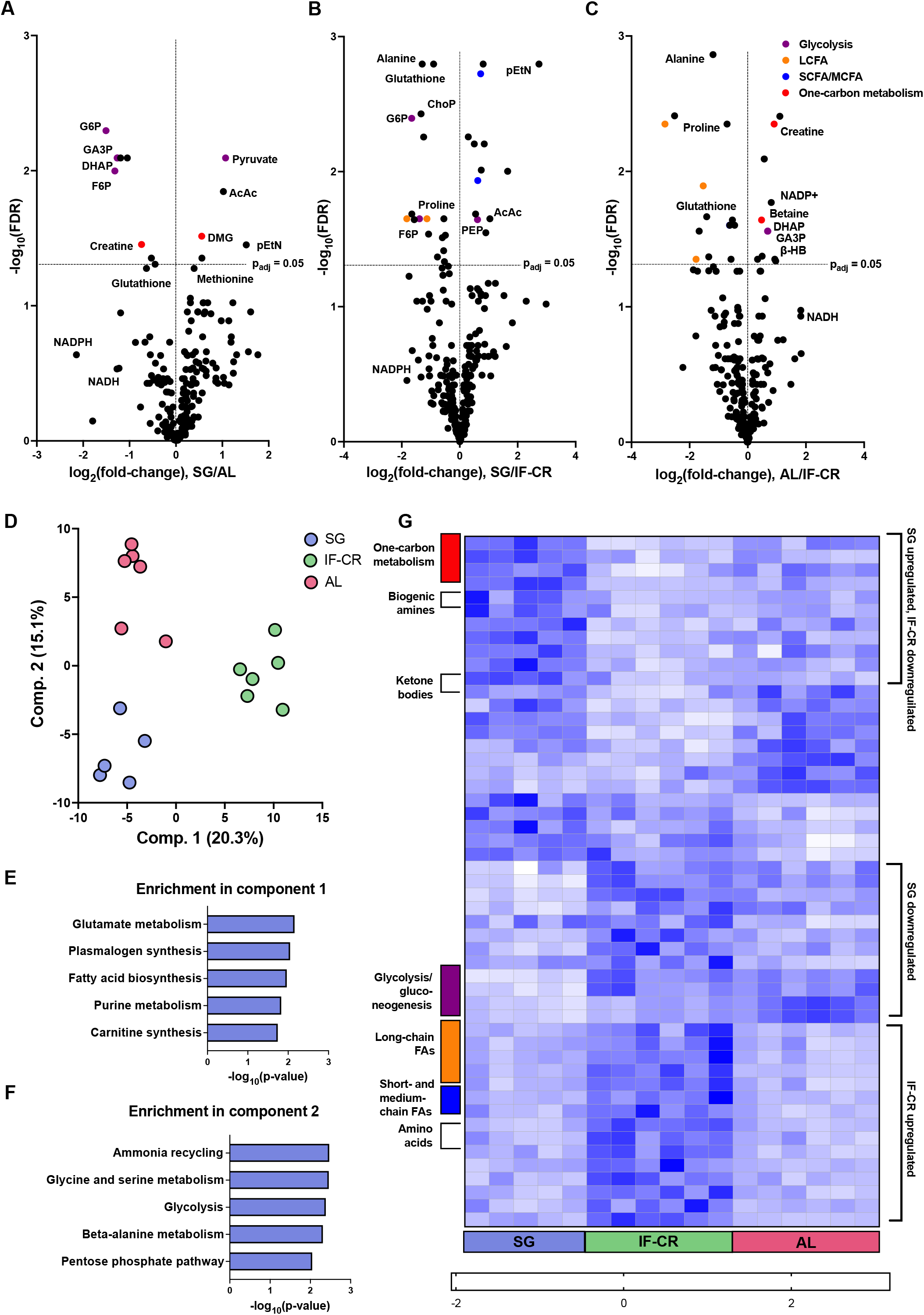
Metabolomic analysis of liver. (A-C) Volcano plots showing the relative changes in metabolite concentrations in the liver between the experimental groups (SG: sleeve gastrectomy; IF-CR: intermittent fasting caloric restriction; AL: ad libitum; FDR = false discovery rate-adjusted p-value of between-group t-tests; LCFA: long-chain fatty acids; SCFA/MCFA: short-/medium-chain fatty acids). (D) Partial least squares-discriminant analysis (PLS-DA) scores plot of metabolite abundance demonstrating separation of the IF-CR group from the AL group along the first component and of the SG group from the AL group along the second component. (E-F) Enrichment analysis by small molecule pathway database (SBPDB) pathways of metabolites in components 1 (E) and 2 (F) of PLS-DA. Metabolites were defined as belonging to component one, two, or neither, based on their VIP score for that component being above 1.0 and greater than the VIP score for the other component. (G) Heatmap of 51 metabolites identified as significantly altered between the experimental groups by ANOVA with FDR of <0.05. Color-code represents log_2_ fold-change between each sample and the mean level of the metabolite.

As in the serum, several metabolites identified by ANOVA showed opposite trends in the SG and IF-CR groups when compared to the AL group (figure 3G). Compounds whose concentrations were increased in the SG groups and decreased in the IF-CR group included one-carbon metabolism-related metabolites and the biogenic amines serotonin and histamine. Showing an opposite pattern were glutathione and the phosphorylated sugars glucose and fructose 6-phosphate. Other metabolites were exclusively affected in the IF-CR group. Consistent with their plasma concentrations, alanine, proline, and isoleucine, and the short-chain fatty acids (butyric and hexanoic acid), were all increased in the IF-CR group. In contrast, the LCFAs, the majority of which showed decreased serum concentrations in the IF-CR group, as mentioned above, were seen to be increased intrahepatically in this group in comparison to the two others, presenting an almost mirror image of that observed in the serum.

### Transcriptomic analysis of liver

2645 differentially expressed genes were identified, of which 1222 were upregulated and 819 were downregulated in the IF-CR group, while 636 were upregulated and 902 were downregulated in the SG group (figure 4A-C). Genes upregulated in the IF-CR group were found to be enriched for citrate cycle, pyruvate metabolism, glycolysis, pentose phosphate pathway, glycine, serine, and threonine metabolism, and fatty acid biosynthesis, while downregulated pathways included fatty acid degradation and linoleic acid metabolism (figure 4D). For the SG group, prominent downregulated pathways were those of glycine, serine, and threonine metabolism, cysteine and methionine metabolism, citrate cycle, and pyruvate metabolism, whereas fatty acid degradation, linoleic acid metabolism, and tryptophan and nicotinamide metabolism were upregulated (figure 4E). PCA showed distinct separation of the IF-CR group from the two other groups along the first principal component, and lesser separation of the SG group from both other groups along the second principal component (figure 4F).

**Figure 4:**
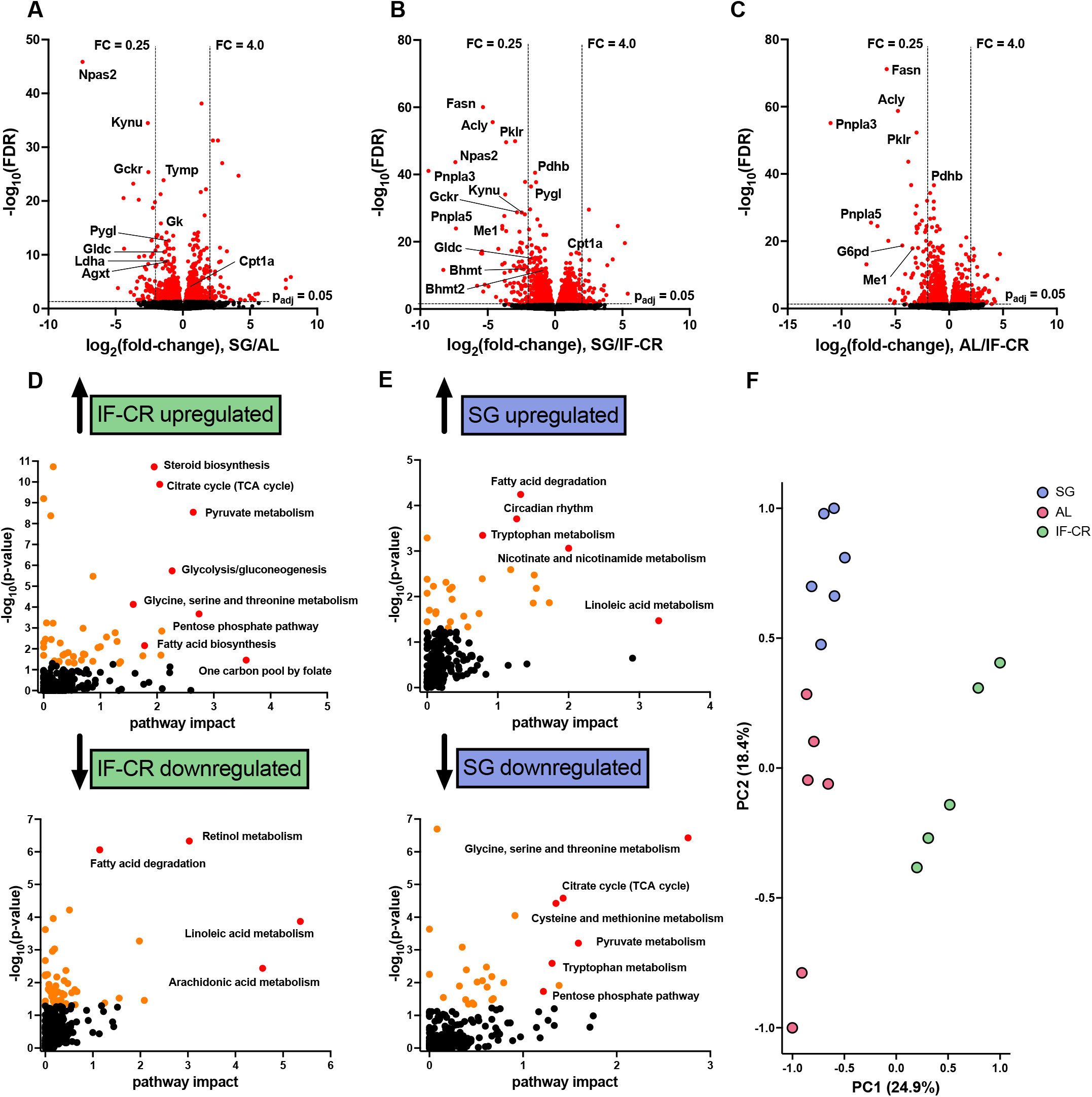
Transcriptomic analysis of liver. (A-C) Volcano plots showing the relative changes in transcript abundance in the liver between the experimental groups (SG: sleeve gastrectomy; IF-CR: intermittent fasting caloric restriction; AL: ad libitum; FDR = false discovery rate-adjusted p-value of between-group t-tests). Red denotes differentially expressed transcripts. (D-E) Enrichment analysis by Kyoto Encyclopedia of Genes and Genomes (KEGG) pathways of genes identified as upregulated (top) or downregulated (bottom) in the IF-CR (D) and SG (E) groups; genes were defined as up- or down-regulated if identified as significantly altered by ANOVA (FDR < 0.05), were identified by post-hoc analysis (Fisher’s least significant difference) as significantly altered against at least one of the other experimental groups, and with either a negative or positive fold-change, respectively, compared to both other groups. (F) Principal component analysis (PCA) scores plot of transcript abundance demonstrating separation of the IF-CR group from the AL group along the first component and of the SG group from the AL group along the second component.

### Fatty acid metabolism: lipogenic pathways increased by IF-CR

In looking jointly at the metabolomic and transcriptomic changes described above, we observe that IF-CR brings about a dramatic change in hepatic lipid metabolism. We witnessed in the IF-CR group a decreased concentration of long-chain FAs (LCFAs) in blood entering the liver but an increased concentration of intrahepatic LCFAs (figure 5A). An almost 3-fold increase in serum triglycerides was seen in the IF-CR group in comparison to the SG group (figure 5B), as well as a decrease in serum ketone bodies (acetoacetate and beta-hydroxybutyrate, figure 5C). These changes were paralleled by an increase in the expression of enzymes involved in DNL and in TG mobilization within the liver in the IF-CR group. In particular, the expression of rate-limiting enzymes of DNL, acetyl-CoA carboxylase (*ACACA*), fatty acid synthetase (*FASN*), acetyl-CoA citrate lyase (*ACLY*), and stearoyl-CoA desaturase (*SCD*)^39^, was increased between 8- and 50-fold in the IF-CR group compared to the AL group. The expression of these enzymes was unchanged in the SG group when compared to the AL group. A concomitant decrease in the expression of key enzymes of fatty acid beta-oxidation was seen in the IF-CR group, including carnitine palmitoyl transferase 1 (*CPT1*) and acyl-CoA oxidase 1 (*ACOX1*), which catalyze the rate-limiting steps in mitochondrial and peroxisomal β-oxidation, respectively^40,41^. These were also seen to be affected by SG, but in an opposite direction, i.e., both were downregulated by approximately 2-fold in the IF-CR group and upregulated by approximately 2-fold in the SG group when compared to the AL-fed animals (figure 5D).

**Figure 5:**
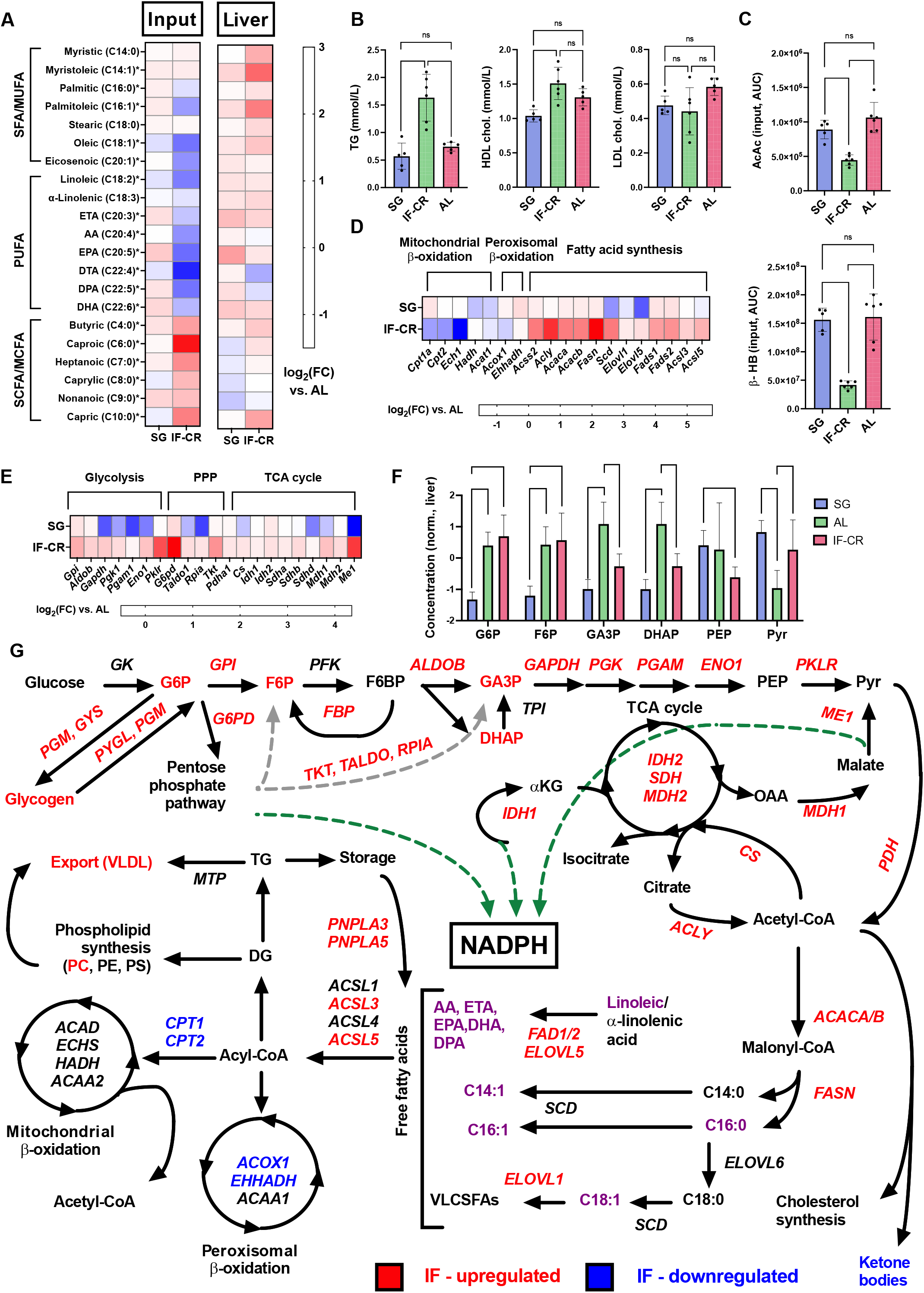
Fatty acid metabolism. (A) Heatmap of concentrations of saturated and mono-unsaturated long-chain fatty acids (SFA/MUFA), poly-unsaturated long-chain fatty acids (PUFA), and short- and medium-chain fatty acids (SCFA/MCFA) in the serum (hepatic input) and liver of the SG and IF-CR groups, expressed as log_2_ of the fold-change versus the AL group. (* - ANOVA FDR < 0.05 in serum or liver). (B) Serum lipid profile showing increased triglyceride (TG) concentrations in the IF-CR group, as well as decreased high-density lipoprotein (HDL) cholesterol levels in the SG group and no change in low-density lipoprotein (LDL) cholesterol levels. (C) Serum concentrations (area under the curve [AUC] normalized to total ion count [TIC]) of the major ketone bodies, acetoacetate (AcAc) and beta-hydroxybutyrate (beta-HB), showing a decrease in the IF-CR group. (E) Heatmaps of relative transcript abundance of significantly altered genes (ANOVA FDR < 0.05) in pathways of fatty acid (FA) oxidation and synthesis, glycolysis, pentose phosphate pathway (PPP), and tricarboxylic acid (TCA) cycle, expressed as log_2_ of the fold-change versus the AL group, and showing upregulation of FA synthesis in the IF-CR group. (F) Liver concentrations of glycolytic intermediates (AUC normalized to TIC, expressed as standardized z-scores) (G6P: glucose 6-phosphate; F6P: fructose 6-phosphate; GA3P: glyceraldehyde 3-phosphate; DHAP: dihydroxyacetone phosphate; PEP: phosphoenol pyruvate; Pyr: pyruvate). (G) plot summarizing changes in fatty acid metabolism pathways, glycolysis, pentose phosphate pathway, and TCA cycle. In red, metabolites or transcripts upregulated in the IF-CR group. In blue, metabolites or transcripts down regulated in the IF-CR group. In purple, long-chain FAs – downregulated in serum and upregulated in the liver of the IF-CR group. * p<0.05 ANOVA with Tukey post-hoc test. ** p<0.01 ANOVA with Tukey post-hoc test.

In parallel with an increase in fatty acid synthetic pathways and in support of lipogenesis, we see in the IF-CR group increased expression of glycolytic, pentose phosphate pathway, and citrate cycle enzymes (figure 5E), and increased levels of glycolytic intermediates in the liver (figure 5F). The changes related to fatty acid metabolism following IF-CR are summarized in figure 5G.

### One-carbon metabolism: decreased utilization of one-carbon donors following SG

As mentioned above, numerous metabolites and transcripts related to one-carbon metabolic pathways were seen to be affected by both SG and IF-CR, in many cases in opposite directions. In the SG group, levels of the major one-carbon donors, including betaine, methionine, serine, glycine, sarcosine, and dimethylglycine (DMG), were increased in both the serum and intrahepatically compared to the AL-fed animals. With IF-CR, decreased levels of betaine, DMG, sarcosine, and glycine, which are intermediates in the sequential breakdown of choline to glycine, were seen in the serum, whereas levels of serine and methionine were increased relative to AL-fed animals (figure 6A). Levels of methyl-utilizing enzymes involved in the transfer of one-carbon groups for regeneration of SAM, including betaine homocysteine S-methyltransferase (*BHMT*), mitochondrial serine hydroxymethyltransferase (*SHMT2*), and glycine decarboxylase (*GLDC*), were all decreased in the SG group compared to the IF-CR group (figure 6B), which, together with the increased levels of methyl donors in this group, suggests reduced consumption of one-carbon groups and decreased transmethylation flux in the post-surgical liver.

**Figure 6:**
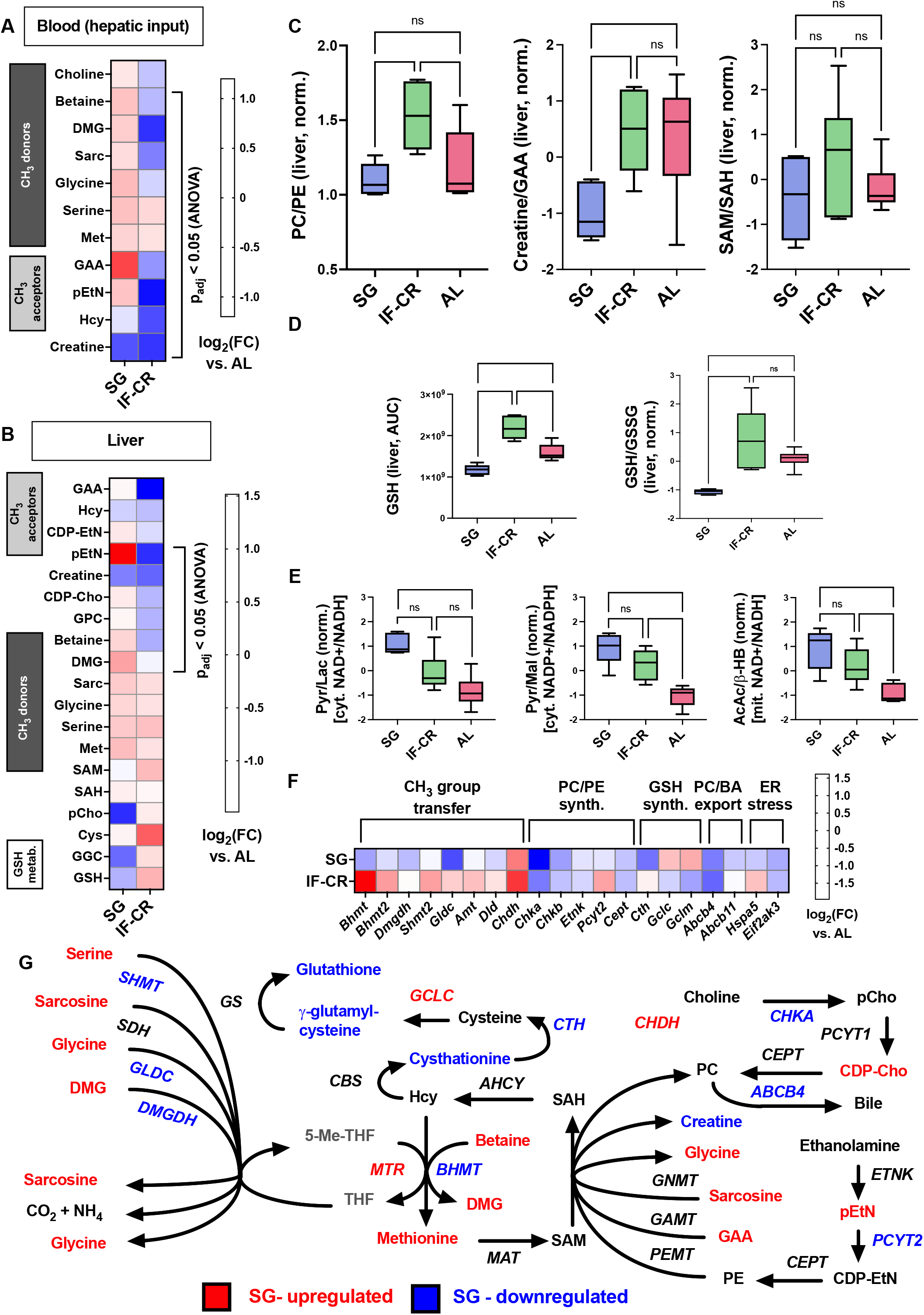
One-carbon metabolism. (A-B) Heatmap of concentrations (area under the curve [AUC] normalized to total ion count [TIC]) of metabolites related to one-carbon transfer in the serum (hepatic input, A) and liver (B) of the SG and IF-CR groups, expressed as log_2_ of the fold-change versus the AL group. (DMG: dimethylglycine; Sarc: sarcosine; Met: methionine; GAA: guanidinoacetate; pEtN: phosphoethanolamine; Hcy: homocysteine; CDP-EtN: cytidine diphosphate-ethanolamine; CDP-Cho: cytidine diphosphate-choline; GPC: glycerophosphocholine; SAM: S-adenosylmethionine; SAH: S-adenosylhomocysteine; pCho: phosphorylcholine; Cys: cystathionine; GGC: gamma-glutamylcysteine; GSH: reduced glutathione). (C) Ratios in liver of methylated phosphatidylcholine (PC) to unmethylated phosphatidylethanolamine (PE), methylated creatine to unmethylated GAA, and SAM to SAH, in the three experimental groups (concentrations in nmol/mg for PC and PE, and AUC normalized to TIC for creatine, GAA, SAM, and SAH, expressed as standardized z-scores). (D) Liver concentrations of reduced glutathione (GSH, AUC normalized to TIC) and ratios of GSH to GSSG (expressed as standardized z-scores). (E) Ratios of hepatic pyruvate to lactate (Pyr/Lac, left), pyruvate to malate (Pyr/Mal, middle), and acetoacetate to beta-hydroxybutyrate (AcAc/beta-HB, right), reflecting cytosolic NAD+/NADH, cytosolic NADP+/NADPH, and mitochondrial NAD+/NADH ratios, respectively (AUC normalized to TIC of each compound was divided by that of the other, and expressed as standardized z-scores). (F) Heatmaps of relative transcript abundance of significantly altered genes (ANOVA FDR < 0.05) related to methyl (CH_3_) group transfer, PC and PE synthesis, GSH synthesis, and PC and bile acid (BA) export. Expressed as log_2_ of the fold-change versus the AL group. (G) Plot summarizing changes in one-carbon metabolic pathways. In red, metabolites or transcripts upregulated in the SG group. In blue, metabolites or transcripts down regulated in the SG group. * p<0.05 ANOVA with Tukey post-hoc test. ** p<0.01 ANOVA with Tukey post-hoc test.

Levels of SAM and SAM/SAH ratios were not significantly altered between the experimental groups (figure 6C). The intrahepatic PC/PE ratio was somewhat increased in the IF-CR group compared to both other groups (figure 6C). Levels of choline kinase (*CHKA*), which catalyzes the rate-limiting step in the synthesis of PC from choline (Kennedy pathway), were reduced post-SG (figure 6F), suggesting that the apparent reduced consumption of methyl groups is not due to increased synthesis of PC via this pathway. Levels of *ABCB4/MDR3*, which is responsible for secretion of PC into bile^42^, were decreased in both the SG and IF-CR groups when compared to the AL group. With regards to the second major methyl-consuming reaction, GAA concentrations were markedly increased post-SG, while the ratio of creatine to GAA was reduced in post-SG livers compared to both control groups (figure 6C), implying decreased hepatic conversion of GAA to creatine in this group as a possible explanation for the reduced utilization of one-carbon groups.

### Redox balance: decreased glutathione concentrations following SG

Consistent with decreased transmethylation activity, we found decreased levels of glutathione in the post-SG livers, as well as a decreased ratio of reduced to oxidized glutathione (GSH/GSSG; figure 6D). This finding contrasts with an increased expression of subunits of the rate-limiting enzyme in GSH biosynthesis, glutamate-cysteine ligase (*GCLC* and *GCLM*; figure 6F), suggesting upstream substrate availability as the cause of decreased GSH post-SG. Additional evidence for a more oxidized metabolic state post-surgery was seen with an increased ratio of acetoacetate to beta-hydroxybutyrate, which provides an estimate of the mitochondrial NAD+/NADH ratio^43^, as well as an increase in the ratios of pyruvate to malate and lactate, which provide an estimate of the cytosolic NADP+/NADPH and NAD+/NADH ratios, respectively (figure 6E). This effect was particularly prominent in comparing the SG to the AL group, and less so with respect to the IF-CR group. Increased glutathione oxidation is linked to attenuated endoplasmic reticulum (ER) stress^44^, and we found decreased expression of unfolded protein response genes (*HSPA5/BIP, EIF2AK3/PERK*), which are markers of ER stress, in the SG group (figure 6F).

The changes in one-carbon and redox pathways following SG are summarized in figure 6G.

## Discussion

In this study, we characterized the changes in liver metabolism following weight loss induced by either SG or IF-CR in a rat model, using combined metabolomic and transcriptomic approaches. We chose an experimental design in which animals were fed a moderately high-fat high-sucrose diet (45% kcal fat) for a relatively short period, to examine the effects of SG and IF-CR in the setting of a high-fat diet but without the metabolic derangements induced by long-term high-fat feeding and hepatic lipid accumulation, and with equal end-experimental weights.

We observed that the two interventions result in significantly different and in some cases opposite effects on hepatic metabolism, despite a similar degree of weight loss. In line with recent findings^7^, IF-CR resulted in changes primarily related to fatty acid metabolism, including marked upregulation of fatty acid synthase (*FASN*) and acetyl-CoA citrate lyase (*ACLY*) and apparent increased TG export. This, together with increased glycogen content, would appear to support energy storage as an adaptation to once-daily feeding.

The metabolic effects of SG were found to be most prominent in one-carbon metabolic pathways. Changes in metabolite and transcript levels following SG point to a decrease in transmethylation flux, as evidenced by an increase in the levels of potential methyl donors and acceptors in the serum. The cause of the decreased utilization of methyl donors post-SG remains unclear. However, increased levels of both methyl donors and methyl acceptors suggest altered regulation at the entry point of the transmethylation cycle itself. That this change may be causative in the metabolic effects seen after bariatric surgery is supported by a recent study showing that silencing of MAT1A prevents obesity and hepatic steatosis^29^. It should be noted that ratios of PC to PE were relatively unaffected following surgery, while GAA to creatine ratios were markedly lowered. Decreased allocation of methyl groups to creatine relative to PC is a pattern also seen with methionine-restricted diets^45^, suggesting functional methionine restriction as can occur with decreased MAT activity, despite an abundance of methionine both in serum and intrahepatically. It should be noted that levels of vitamin B12, an essential cofactor in folate-mediated remethylation of homocysteine, have been known to decrease in some cases following bariatric surgery^46^. However, development of such a deficiency has been shown to result in compensatory overutilization of choline and betaine as methyl donors^47^, and thus would be expected to result in depletion of these metabolites, rather than increased levels as seen in our study.

Our results are in agreement with a recent systematic review on the metabolic changes associated with bariatric surgery in general and a recent study examining the effects of gastric bypass on the metabolic composition of the portal blood, both of which identified glycine, serine, and threonine metabolism and cysteine and methionine metabolism as the metabolic pathways uniquely affected by bariatric surgery^48,49^. A decrease in SAM-synthesizing enzymes was described in mice following duodenal-jejunal bypass (DJB)^50^. A similar profile of increased levels of methyl donors and acceptors as seen in our study was observed in humans following Roux-en-Y gastric bypass, where levels of both methionine and PE were found to inversely correlated with insulin resistance following surgery^51^.

Levels of glycine have been consistently observed to be lower in individuals with obesity and the metabolic syndrome, and to increase following bariatric surgery^52^. While other explanations for this phenomenon have been proposed^53^, our study suggests decreased hepatic glycine catabolism, reflecting a decreased demand for methyl donors, as a possible cause of increased circulating glycine levels following bariatric surgery.

Alterations in transmethylation in the SG group in our experiment also affected the interlinked transsulfuration cycle, resulting in an apparent decrease in hepatic antioxidant defenses. MAT activity is known to be reduced by oxidative stress, an effect which appears paradoxical but has been demonstrated in multiple studies^54–56^. Although transcript levels of the various MATs were not different between the experimental groups, the alterations in redox metabolism may underlie what appears as a reduced entry of methionine into the transmethylation cycle. While oxidative stress is generally thought of as a deleterious phenomenon, the early stages of the development of liver steatosis are characterized by increased antioxidant capacity and GSH synthesis^57,58^, and several studies have pointed to a beneficial effect of reactive oxygen species (ROS) generation and glutathione depletion early in the development of NAFL^29,44,59–61^ or in the context of a methionine- and choline-deficient diet^61^. Indeed, so-called reductive stress is now appreciated to be in some cases as harmful as excess oxidative stress and a direct cause of ER stress^62^. Transcript levels of proteins involved in the ER stress response were lower post-SG in our study. A decrease in antioxidant capacity may underlie both the beneficial effect in the majority of patients, and the puzzling worsening of NAFLD features in a subset (approximately 10%) of patients undergoing bariatric surgery^63^.Decreased concentrations of serum GSH following bariatric surgery and an increased susceptibility to acetaminophen toxicity have been reported in several small human studies^64–66^, although others have reported an opposite effect^67^. A single animal study evaluating metabolic changes in the liver tissue itself found increased rather than decreased GSH levels after bariatric surgery, in apparent direct opposition to our results^68^; however, this may be due to the different surgical interventions undertaken (DJB vs. SG) or to different degrees of weight loss induced by these interventions, as the animals that underwent DJB experienced a gain in body weight of nearly 150% from their pre-surgery level over a period of 5 weeks, compared to a 5-10% increase in body weight over 4.5 weeks in our study.

Our study has several limitations. First, we obtained a snapshot view of metabolite and transcript levels at a certain time of day and after a predefined period of fasting. Therefore, we are unable to distinguish whether the observed alterations in metabolite levels are due to changes in absorption, synthesis, or consumption, despite some circumstantial evidence pointing towards one of these options in the case of specific metabolites, such as an increase in the levels of serum betaine paralleled by a decrease in the levels of enzymes involved in betaine catabolism in the liver. This may be validated in the future by metabolic flux studies utilizing stable isotope labeling of metabolites of interest. A further limitation brought about by the study design is that between-group differences may be restricted to the specific time point chosen, rather than reflect a generally altered metabolic state.

In conclusion, our study highlights the disparate metabolic effects of different weight-loss interventions (figure 7). Surprisingly, the two weight loss interventions shared only few features from the standpoint of changes in hepatic metabolism, despite a similar degree of weight loss. While IF-CR resulted in sweeping changes across multiple central metabolic pathways, the effects of SG were seen to be more modest and concentrated on one-carbon metabolic pathways which may affect redox balance. Further studies are required to determine the mechanistic link between IF-CR and particularly SG on hepatic metabolism, and to understand post-prandial hepatic metabolism following these interventions.

**Figure 7:**
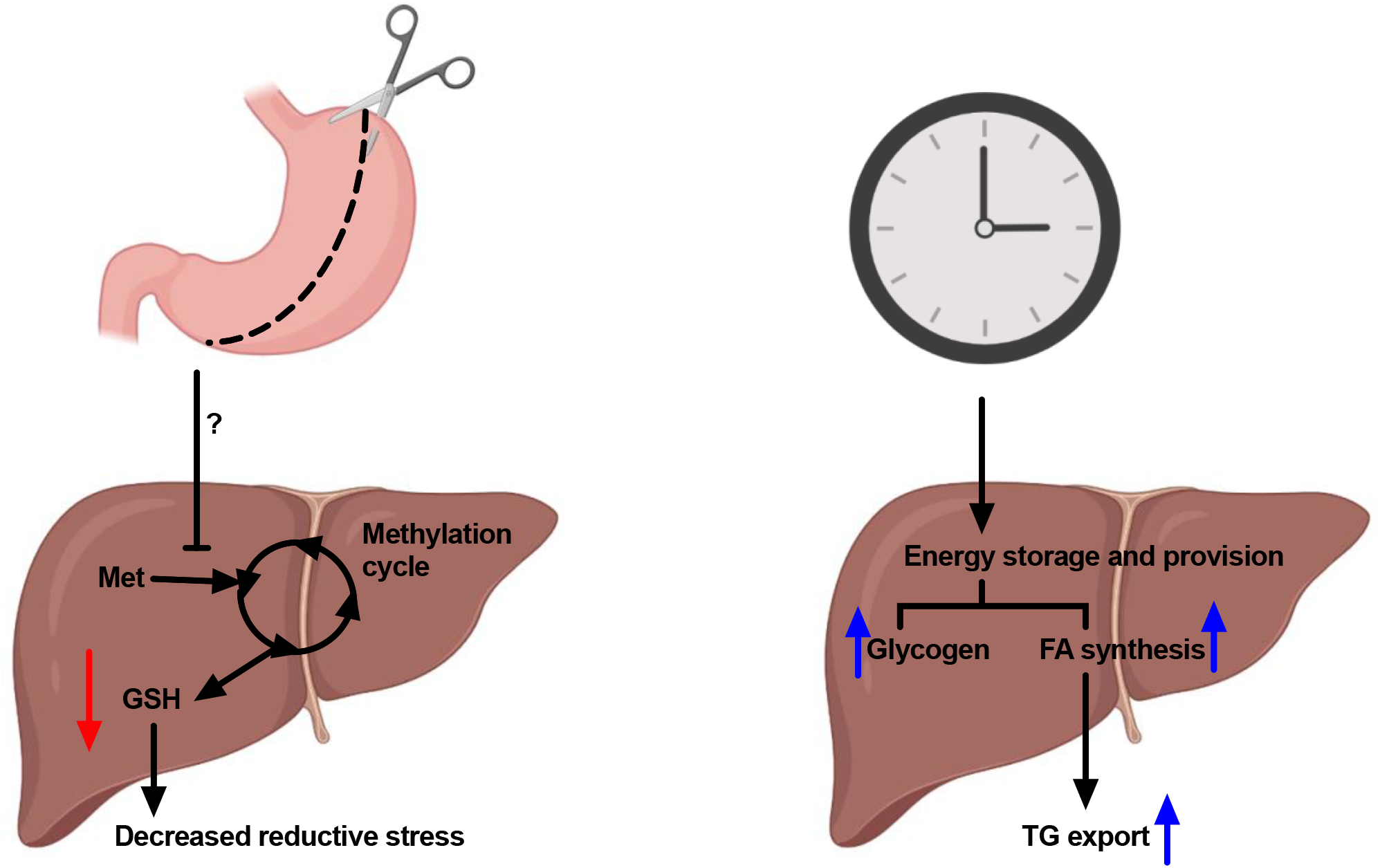
The effects of weight loss through bariatric surgery and intermittent fasting with caloric restriction on hepatic metabolism.

